# Variable Cerebral Blood Flow Responsiveness to Acute Hypoxic Hypoxia

**DOI:** 10.1101/2024.11.14.623430

**Authors:** Hannah R Johnson, Max C Wang, Rachael C Stickland, Yufen Chen, Todd B Parrish, Farzaneh A Sorond, Molly G Bright

## Abstract

Cerebrovascular reactivity (CVR) to changes in blood carbon dioxide and oxygen levels is a robust indicator of vascular health. Although CVR is typically assessed with hypercapnia, the interplay between carbon dioxide and oxygen, and their ultimate roles in dictating vascular tone, can vary with pathology. Methods to characterize vasoreactivity to oxygen changes, particularly hypoxia, would provide important complementary information to established hypercapnia techniques. However, existing methods to study hypoxic CVR, typically with arterial spin labeling (ASL) MRI, demonstrate high variability and paradoxical responses. To understand whether these responses are real or due to methodological confounds of ASL, we used phase-contrast MRI to quantify whole-brain blood flow in 21 participants during baseline, hypoxic, and hypercapnic respiratory states in three scan sessions. Hypoxic CVR reliability was poor-to-moderate (ICC=0.42 for CVR relative to P_ET_O_2_ changes, ICC=0.56 relative to SpO_2_ changes) and was less reliable than hypercapnic CVR (ICC=0.67). Without the uncertainty from ASL-related confounds, we still observed paradoxical responses at each timepoint. Concurrent changes in blood carbon dioxide levels did not account for paradoxical responses. Hypoxic CVR and hypercapnic CVR shared approximately 40% of variance across the dataset, indicating that the two effects may indeed reflect distinct, complementary elements of vascular regulation.

## Introduction

Cerebral blood flow (CBF) is regulated to ensure that sufficient oxygen and nutrients are delivered to brain tissue despite changes in metabolic demands, blood pressure, and blood gas levels. The ability of the brain vasculature to appropriately regulate blood flow can be assessed by measuring cerebrovascular reactivity (CVR), the change in CBF in response to a vasoactive stimulus like hypercapnia, hypoxia, or acetazolamide^1^. CVR is a useful metric of vascular health, changing with age^2^ and decreasing with disease severity in Alzheimer’s Disease^3^, Multiple Sclerosis^4^, Parkinson’s Disease^5^, and Obstructive Sleep Apnea^6^. CVR is also a promising predictive biomarker of stroke in carotid artery disease^7,8^ and white matter hyperintensity development in white matter disease^9^, increasing its clinical utility.

CVR is most commonly assessed using a hypercapnic stimulus, such as a breath-holding task or inhalation of gas mixtures with elevated CO_2_, as in the aforementioned studies. CVR to hypercapnic stimuli has been extensively characterized, with a large body of literature establishing normative values for multiple imaging modalities and evaluating the impact of a wide range of clinical conditions on CVR^1,10^. Although the cerebrovascular response to hypercapnia certainly provides valuable information about the health of the brain’s blood supply, the brain is also exquisitely sensitive to hypoxic conditions due to the physiological importance of oxygen^11^. While oxygen and carbon dioxide levels are closely coupled through ventilation under healthy conditions, their relationship can change under pathology, particularly in respiratory disease and ischemia^12–14^, and the two influence vascular tone through independent mechanisms^15,16^. Characterization of the hypoxic vascular response, independent of carbon dioxide changes, could provide complementary information to the extensive body of literature on the hypercapnic cerebrovascular response and inform assessment of composite CVR in pathology.

There are technical challenges to studying CVR to hypoxia, however. Spontaneous breathing during hypoxic hypoxia (poikilocapnic conditions) results in decreased arterial CO_2_ concentrations, producing a vasoconstrictive effect that competes with and obscures the anticipated hypoxic vasodilatory response we seek to characterize^17,18^. While transcranial doppler (TCD) provides unbiased measures of blood velocity changes in hypoxia, it is unable to account for arterial diameter changes and therefore cannot fully characterize CVR^19^. Hypoxia also introduces bias in typical MRI methods for assessing CVR: bulk changes in arterial oxygen confound characterization of CBF changes with blood oxygenation level-dependent (BOLD)-weighted functional MR imaging, and changes in blood velocity and arterial oxygenation can impact measurements of quantitative CBF with arterial spin labeling (ASL) MRI. Although the spatial distribution of CBF changes is lost, phase-contrast (PC) MRI offers a robust method for quantifying whole-brain CBF changes during hypoxia, accounting for modulations in both velocity and vessel area, and unbiased by BOLD-weighted contrast or ASL confounds related to labeling efficiency and T1 relaxation times.

Considering all imaging methods, the blood-flow response to severe hypoxia has been well-characterized, causing an invariable exponential increase in CBF with decreases in the partial pressure of oxygen^11,20,21^, but the response to mild-to-moderate hypoxia, particularly when assessed with non-invasive methods, remains ambiguous. While there is no current standardized classification system for defining hypoxia as mild, moderate, or severe, general thresholds exist^22^. Mild hypoxia generally delivers a fraction of inspired oxygen (FiO_2_) of greater than 17% but less than room air, corresponding to inhaled partial pressures of oxygen (PO_2_) between 130 and 160 mmHg at sea level. Moderate hypoxia delivers FiO_2_ of between 10 and 16%, corresponding to PO_2_ of 76-130 mmHg. Severe hypoxia delivers air with less than 9% oxygen, equating to PO_2_ of less than 76 mmHg. As the typical PaO_2_/FiO_2_ ratio for healthy controls at sea level is between 400 and 500 mmHg^23^, these thresholds approximately correspond to PaO_2_ levels above 75 mmHg and SpO_2_ levels above 94% for mild hypoxia, PaO_2_ between 40 and 75 and SpO_2_ between 75-94% for moderate hypoxia, and PaO_2_ below 40 and SpO2 below 75% for severe hypoxia in healthy individuals^24^.

At the (generally safe) levels of oxygen delivery used in mild-to-moderate hypoxia challenges, the blood flow response is comparatively small, limiting its detection, and has shown substantial variability between individuals. The typical response to mild-to-moderate hypoxia is an increase in blood flow. Kety and Schmidt first measured this response using arterial sampling, finding an average CBF increase of 37% (±23%) in response to 10% inhaled oxygen^25^. Later, Ellingsen et al. found an invariable increase in blood velocity in the internal carotid artery using doppler ultrasound, averaging a 23% (range 12.5-25%) increase in CBF, when inhaled oxygen was reduced by 35% to approximately 13.7%^26^. More recently, a 19% (±3%) increase in total CBF during an 11% inhaled oxygen challenge was measured using 4D-flow MRI^27^. However, several published studies have shown that a subset of participants deviate from this expected response, exhibiting either no significant change or a paradoxical decrease in CBF during hypoxia administration. In an ASL MRI study, 30% of participants exhibited decreased CBF to moderate hypoxia, with an average decrease in blood flow of .68% (±.31%) per % change in SaO_2_^28^. In another study using positron emission tomography, Binks et al. observed that one of five participants had no change in CBF under moderate hypoxia^14^. These studies demonstrate considerable deviation from the expected blood flow response to hypoxia, and such variable responsiveness could cause some individuals to have greater susceptibility to cerebral tissue deoxygenation. However, it is unclear whether this responsiveness to hypoxia is an artifact of methodological confounds, a result of day-to-day variability in physiology, or is a true representation of vascular reactivity in some individuals.

We therefore used phase-contrast MRI, which is free of the biases associated with other techniques, to quantify whole-brain blood flow under baseline, hypoxic, and hypercapnic respiratory conditions on three days spanning several months. We characterized inter-subject and inter-session variability in the cerebrovascular response to moderate hypoxic hypoxia while accounting for important sources of physiological variability. We further compared hypoxic CVR with hypercapnic CVR measures in the same participants, evaluating the differences between the two metrics of vascular reactivity. Through this robust evaluation of CVR to mild-to-moderate hypoxia in healthy individuals, we hope to improve the utility of hypoxic CVR as a metric of cerebrovascular health and characterize sources of variability.

## Materials and Methods

This study was approved by Northwestern University’s Institutional Review Board. All subjects provided written informed consent before undergoing any study procedures. 21 subjects (8 male, 12 female, 1 non-binary, 28±7 years) were recruited. Subjects were free of known neurological, vascular, or respiratory conditions, had no MRI contraindications, and were not pregnant. Participants were scanned twice, 21±5 days apart, following an initial practice session outside of the scanner to ensure safety and comfort with gas challenges. Scans were scheduled at approximately the same time of day to minimize diurnal variations in blood flow and CVR, and subjects were asked to refrain from caffeine consumption for at least 2 hours before scanning. 19 participants were scanned a third time, 124±29 days (approximately 4 months) after the first MRI session using the same protocol. Note that scans were part of a larger randomized cross-over clinical trial with washout evaluating the effect of a respiratory intervention on CBF and CVR metrics; this study uses data from the sham intervention arm and from the first scan of the intervention arm but excludes data affected by the intervention.

### Gas Protocol

Subjects were fitted with an airtight face mask for scanning and connected to a computer-controlled gas blending system (RespirAct, Thornhill Medical, Toronto, Canada) used to prospectively target precise end-tidal partial pressures of oxygen (P_ET_O_2_) and carbon dioxide (P_ET_CO_2_) through manipulation of inhaled gas concentrations. The respiratory protocol consisted of 4 phases: (1) baseline, targeting participants’ resting P_ET_O_2_ and P_ET_CO_2_ levels, (2) hypoxia, targeting P_ET_O_2_ of 55-70mmHg and resting P_ET_CO_2_, (3) recovery baseline, targeting resting P_ET_O_2_ and P_ET_CO_2_, and (4) hypercapnia, targeting resting P_ET_O_2_ and P_ET_CO_2_ of 6mmHg above resting. The respiratory stimulus protocol is shown in Figure 1. Appropriate targets for P_ET_O_2_ during hypoxia blocks, intended to maintain an oxygen saturation of at least 85%, were determined on a subject-specific basis during the practice session and ranged between 55 and 70mmHg. Dynamic achieved P_ET_O_2_ and P_ET_CO_2_ were recorded continuously for the entire scan session through the gas blending system alongside respiratory rate and tidal volume. Participant SpO_2_ and heart rate were monitored continuously throughout scanning via a finger clip sensor, and values were recorded at the start of each PC scan. Blood pressure was taken before and after scanning.

**Figure 1.**
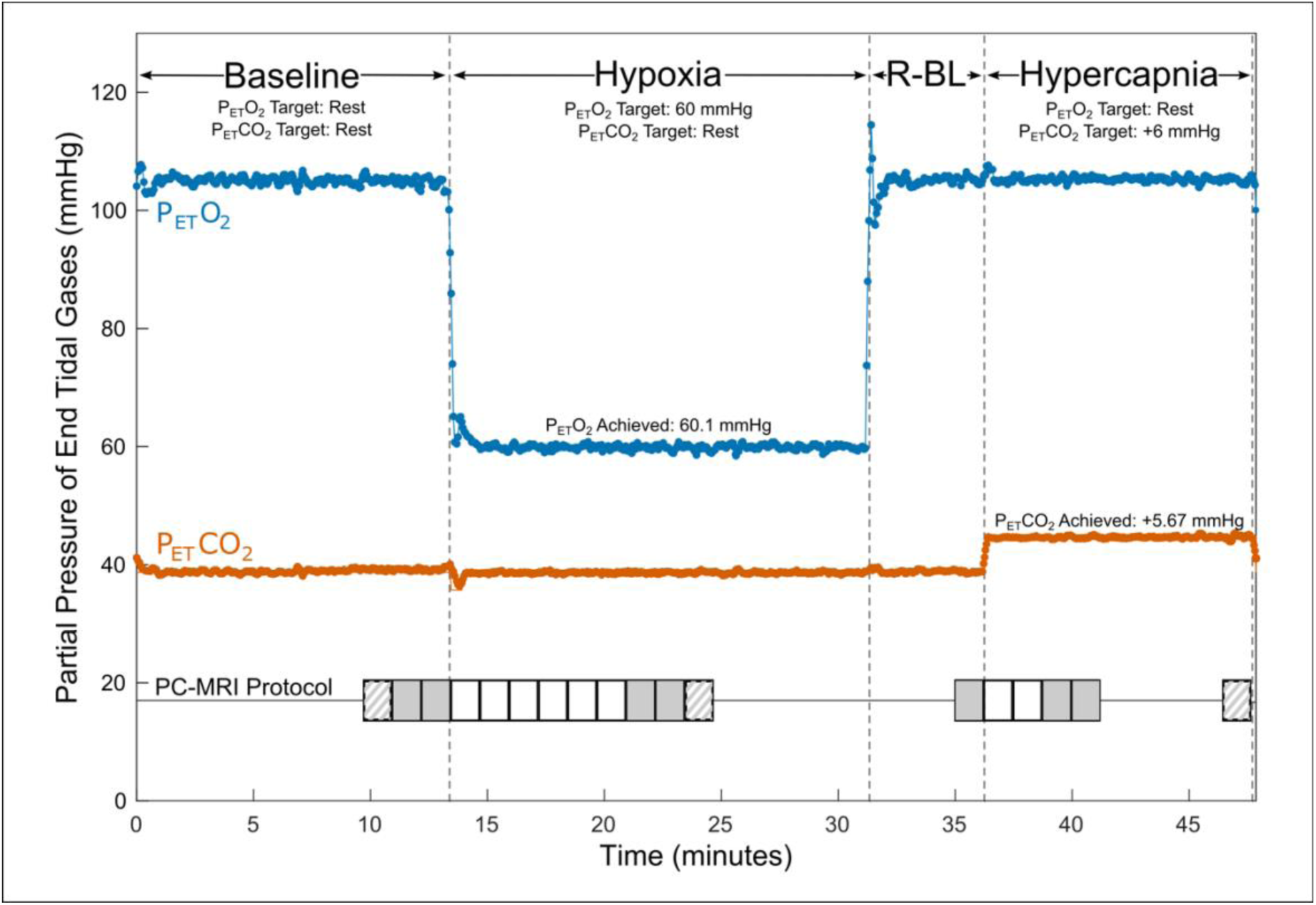
Respiratory and PC-MRI Acquisition Protocol. P_ET_CO_2_ (shown in orange) and P_ET_O_2_ (shown in blue) values are shown for an example participant during baseline, hypoxia, recovery baseline (R-BL), and hypercapnia. Baseline targeting lasted a minimum of 12 minutes (median 18 minutes), hypoxic targeting lasted at least 13 minutes (median 16 minutes), recovery baseline targeting lasted 5 minutes, and hypercapnic targeting lasted a minimum of 9 minutes (median 12 minutes). Timings of PC acquisitions are shown relative to respiratory state. Solid grey shading indicates placement 1 acquisitions used in calculations of CVR. Striped shading indicates placement 2 acquisitions that may have been used in calculations of CVR. Solid white shading indicates placement 1 acquisitions that were not used in CVR calculations. Additional scanning was completed between PC acquisitions.

### Scanning

Scanning was performed on a Siemens 3T Prisma scanner; a 32-channel head coil was used to allow space for the face mask. A 3D time-of-flight angiogram (TR=21.0ms, TE=3.41ms, voxel size=0.8×0.8×1.2mm^3^) was used to manually position imaging slices for subsequent PC measures. Slices were chosen perpendicular to the internal carotid and vertebral arteries, superior to the carotid bifurcation. Two placements were selected: placement 1 prioritized orthogonality of the imaging slice with the carotid arteries, and placement 2 prioritized orthogonality with any of the four vessels that were not considered orthogonal in placement 1, typically one or both vertebral arteries. These placements are shown for an example participant in Supplemental Figure 1. Slice positioning was replicated in subsequent scan sessions using screenshots from the first session. Consecutive retrospectively gated PC scans (TR=47.52ms, TE=3.45ms, voxel size=0.5×0.5×5.0mm^3^, encoding velocity=100cm/s, flip angle=15°, 8 frames per cardiac cycle) were acquired thrice (once at placement 2, twice at placement 1) during baseline gas delivery and repeatedly during the transition to steady-state, with three additional scans (two at placement 1, one at placement 2) collected at estimated steady-state hypoxia. Previous literature suggests that approximately 7.5 minutes are needed for blood flow to stabilize following the onset of hypoxia^21^; based on this and pilot data, we allowed approximately 7.5 minutes before collecting steady-state scans. PC images were also collected once at the end of the recovery baseline, repeatedly during the transition to steady-state hypercapnia, and thrice at steady-state hypercapnia, allowing approximately 2.5 minutes for blood flow to stabilize. Images collected during the transitions to steady-state were all acquired at placement 1 (Fig. 1). A T1-weighted MPRAGE scan (TR=2170.0ms, TE=1.69/3.55/5.41ms, voxel size=1.0×1.0×1.0mm^3^, FOV read=256mm, flip angle=7.0°) was acquired during baseline for calculation of brain volume.

### Data and Statistical Analysis

The T1 image collected in session 1 was processed with FSL-BET and FAST for brain extraction and segmentation^29,30^. Grey matter and white matter volumes were summed to calculate total brain volume for each participant. Total brain mass was calculated from the total brain volume, assuming a tissue density of 1.06 g/mL^31,32^.

CBF was calculated for each PC-MRI acquisition using cvi42’s Flow 2D (Circle Cardiovascular Imaging, Calgary, Canada). Images were corrected for background phase errors. Regions of interest for internal carotid and vertebral arteries were manually selected for each cardiac bin in each acquisition. Mean flow, the product of average velocity and cross-sectional area, across the cardiac cycle for each vessel was calculated. The mean flows of the four vessels were summed to find total CBF in units of mL/min and further divided by the brain mass to find normalized CBF (nCBF), with units of mL/100g/min.

Each reported measurement of CBF combines measurements from two or three acquisitions. Carotid artery blood flow measurements are taken as the average of the two steady-state PC acquisitions at placement 1. For vertebral arteries, the relationship between steady-state blood flow captured by placements 1 and 2 was modeled and used to identify outliers from this relationship, indicating that flow through the artery was not well-captured by placement 1. If the flow measured by placement 2 exceeded the flow measured by placement 1 by greater than 1.96 times the standard deviation of the sum-squared errors of the modeled group relationship at any point, the placement 2 flow values for that vertebral artery were used to calculate total CBF for all timepoints within each session rather than the average of the placement 1 flow values. Recall that placement 2 was generally chosen to better optimize orthogonality with one or both vertebral arteries and is therefore expected to give higher, and ostensibly more accurate, estimates of vertebral blood flow when placement 1 is inadequate.

End-tidal data were temporally aligned with scan acquisitions. A start marker was inserted in respiratory data synchronized with the start of the first baseline PC acquisition during data collection. The start time, relative to this first marked scan, and the length of each PC acquisition were calculated from header data and used to find the start and end time of each acquisition relative to the continuous respiratory data; the average P_ET_CO_2_, P_ET_O_2_, and minute ventilation across each PC-MRI acquisition was then calculated. For simplicity, average end-tidal and physiological data from placement 1 acquisitions were used for all comparisons and calculations, even when select vessel data were taken from placement 2 as described above. We make the assumption that this difference is small, given that measures are taken under identical physiological conditions and the time between measurements is on the order of 1-2 minutes.

CVR was calculated for hypoxic and hypercapnic conditions by finding the change in CBF as a function of the change in end-tidal partial pressure (Equations 1 and 2) or as a function of the change in oxygen saturation (Equation 3). Data from the initial baseline, rather than recovery baseline, were used for all calculations.

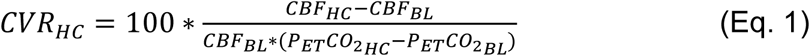

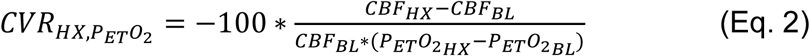

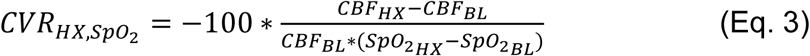

The mean and standard deviation of end-tidal gases, SpO_2_, heart rate, minute ventilation, and nCBF were calculated for each experimental phase; we tested for significant differences with a mixed-effects model accounting for subjects and sessions as random effects and post-hoc testing to compare stimuli to baseline. We verified that steady-state blood flow was sufficiently achieved through paired two-sided Wilcoxon tests of the differences in back-to-back CBF measurements at placement 1, considering repeated measures during baseline, hypoxic, and hypercapnic experimental phases. Furthermore, immediate test-retest repeatability from these back-to-back measures was assessed with the intra-class correlation coefficient (ICC).

Paired two-sided Wilcoxon tests were used to verify that the distributions of CBF changes due to stimuli were different than the distribution of error in CBF measurements during baseline. The reliability of CVR metrics across sessions was assessed using ICC and standard deviation (SD) estimates. ICCs were calculated in R Version 4.3.2 with software package ‘irr’ Version 0.84, with a one-way random effects model for absolute agreement between the averaged CVR metrics for each session and respiratory stimulus in complete datasets. Inter-subject and inter-session SDs were derived from a linear mixed-effects model, considering only complete datasets, with subject included as a random effect. Coefficients of determination were calculated for the relationships in hypoxic CVR and hypercapnic CVR between sessions, and between the two stimuli.

## Results

Nineteen subjects completed all three scan sessions; two subjects were unable to return for the third scan session. The hypercapnic portion of the scan was not completed in five otherwise complete scan sessions due to time constraints, and SpO_2_ data were not recorded in Session 1 for Subject 1. Details of participant data inclusion, as well as calculated CVR values, are summarized in Supplemental Table 1. Summary metrics of blood flow, gas levels, and physiological measures during each experimental phase are shown in Table 1, calculated from all available data. At the group level, during hypoxia, P_ET_O_2_ and SpO_2_ decreased significantly compared to baseline, and heart rate and minute ventilation increased significantly. P_ET_CO_2_ increased significantly during hypercapnia, as did minute ventilation. No significant differences were present between the two phases of baseline targeting.

**Table 1.**
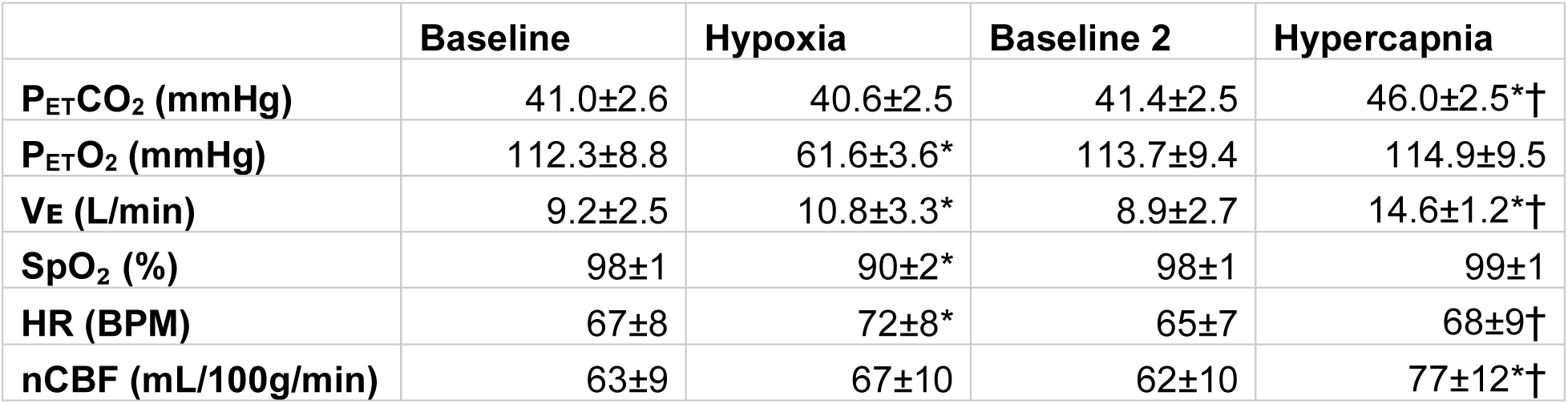
Achieved CBF, P_ET_CO_2_, P_ET_O_2_, and physiological parameters during each respiratory state. Group mean values across all scan sessions during steady-state PC acquisitions. * indicates a significant difference (p less than a threshold of 0.05/18 = 0.00278) compared with BL1; † indicates a significant difference compared with BL2 for hypercapnia. Note that P_ET_CO_2_, P_ET_O_2_, and minute ventilation (V_E_) metrics reported are averaged across the acquisition time; heart rate (HR) and oxygenation (SpO_2_) values were recorded at the beginning of each acquisition.

|  | Baseline | Hypoxia | Baseline 2 | Hypercapnia |
| --- | --- | --- | --- | --- |
| P<sub>ET</sub>CO<sub>2</sub> (mmHg) | 41.0±2.6 | 40.6±2.5 | 41.4±2.5 | 46.0±2.5*† |
| P<sub>ET</sub>O<sub>2</sub> (mmHg) | 112.3±8.8 | 61.6±3.6* | 113.7±9.4 | 114.9±9.5 |
| V<sub>E</sub> (L/min) | 9.2±2.5 | 10.8±3.3* | 8.9±2.7 | 14.6±1.2*† |
| SpO<sub>2</sub> (%) | 98±1 | 90±2* | 98±1 | 99±1 |
| HR (BPM) | 67±8 | 72±8* | 65±7 | 68±9† |
| nCBF (mL/100g/min) | 63±9 | 67±10 | 62±10 | 77±12*† |

The immediate repeatability of CBF metrics was good, with back-to-back baseline measurements at placement 1 having an ICC of 0.88. The differences in CBF between back-to-back PC measurements during hypoxia or hypercapnia were not significantly different from differences in back-to-back PC measurements during baseline, indicating that steady-state blood flow was sufficiently reached (Supplemental Figure 2). Overall, back-to-back measurements showed a median absolute difference of 2mL/100g/min; this difference reflects the inherent influence of both measurement noise and natural physiological fluctuations on CBF measures. Consequently, in this study, we consider changes in CBF exceeding this threshold to be meaningful. The distributions of hypoxic nCBF changes (p<0.001) and hypercapnic nCBF changes (p<0.001) were significantly different from the null distribution of back-to-back changes in nCBF.

Mean total CBF during baseline was 779 mL/min, corresponding to a nCBF of 63 mL/100g/min. CBF increased during hypoxia, with an average increase of 3±4 mL/100g/min or 5±7%. CBF also increased during hypercapnia, with an average increase of 14±7 mL/100g/min or 22±11%, shown in Figure 2. Percent changes are shown in Supplementary Figure 3. The average hypoxic CVR for the group was 0.10±0.14%/- mmHg P_ET_O_2_ and 0.58±0.88%/-%SpO_2_. Average hypercapnic CVR was 4.33±1.83%/mmHg P_ET_CO_2_.

**Figure 2.**
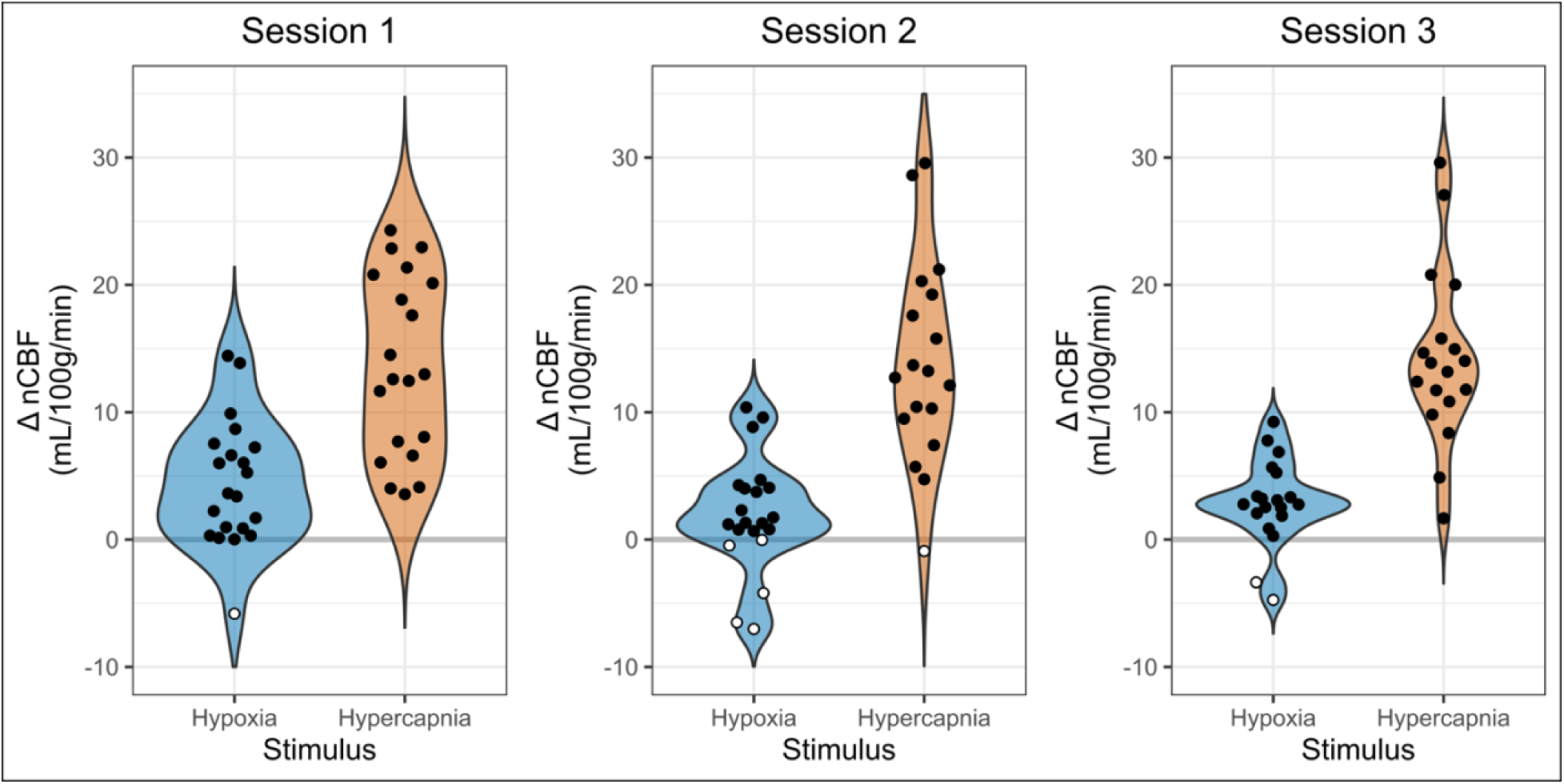
Changes in Blood Flow During Hypoxia and Hypercapnia. Changes in nCBF during hypoxia (blue) and hypercapnia (orange), relative to baseline, are shown for each session. Positive values, indicating an increase in CBF, are shown with black circles, while negative values, indicating a decrease in CBF, are shown with white circles. Note that sessions and gas stimuli have different numbers of subjects; all available data are shown.

### CVR Reliability

The reliability of CVR measures is shown in Figure 3. Only subjects with complete datasets, Group A, were included in overall reliability metrics considering all three timepoints (see Supplemental Table 1 for individual CVR metrics and which subjects were included in groups). Hypoxic CVR showed relatively greater variability between subjects and between sessions than hypercapnic CVR, and inter-session variability dominated. Hypoxic CVR relative to changes in P_ET_O_2_ showed an inter-subject standard deviation (SD) of 0.07%/-mmHg P_ET_O_2_ and an inter-session SD of 0.14%/-mmHg P_ET_O_2_; hypoxic CVR relative to changes in SpO_2_ showed slightly greater variability due to subject, with an inter-subject SD of 0.48%/-%SpO_2_ and an inter-session SD of 0.75%/-%SpO_2_. Hypoxic CVR was found to have overall ICCs of 0.42 and 0.56 relative to changes in P_ET_O_2_ and SpO_2_, respectively. Hypercapnic CVR demonstrated substantially less variability relative to the magnitude, with an inter-subject SD of 1.13%/mmHg P_ET_CO_2_, an inter-session SD of 1.37%/mmHg P_ET_CO_2_, and an ICC of 0.67.

**Figure 3.**
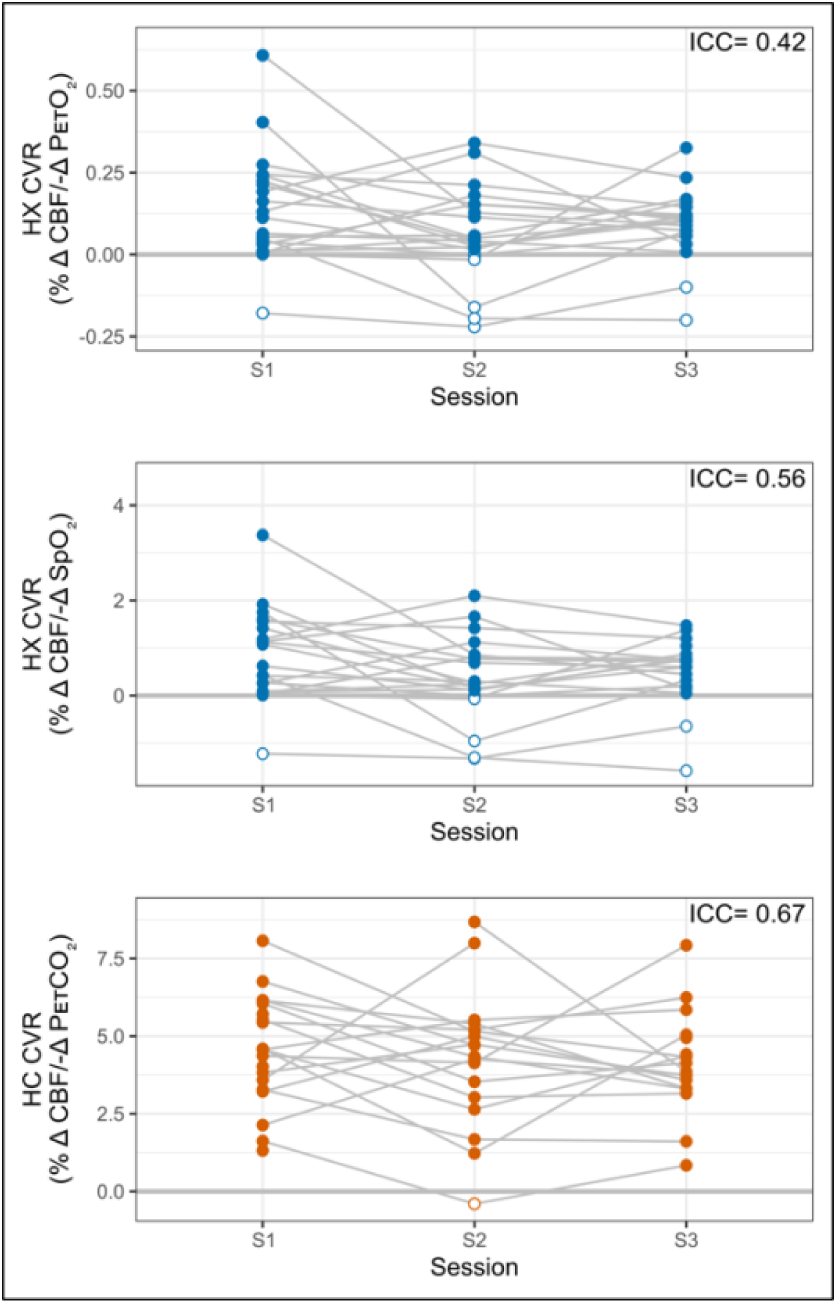
Reliability of Hypoxic and Hypercapnic CVR Measures. Hypoxic (HX) CVR metrics across sessions are shown, relative to P_ET_O_2_ (top) and SpO_2_ (middle). Hypercapnic (HC) CVR is shown, relative to P_ET_CO_2_, for comparison (bottom). Overall intraclass correlation coefficients (ICCs) are listed for subjects with complete datasets (Group A). Negative CVR values are indicated with open circles; all available data are shown.

ICCs considering only sessions 1 and 2, representing reliability over 3 weeks, were 0.33 for P_ET_O_2_ hypoxic CVR, 0.50 for SpO_2_ hypoxic CVR, and 0.57 for hypercapnic CVR. Reliability for all CVR metrics decreased with time: ICCs considering only sessions 1 and 3, representing reliability over 3-5 months, were 0.20 for P_ET_O_2_ hypoxic CVR, 0.23 for SpO_2_ hypoxic CVR, and 0.48 for hypercapnic CVR.

### Minimal and Negative Changes in CBF during Hypoxia

Hypoxia is expected to produce a positive change in CBF. However, previous literature in hypoxia reports observations of paradoxical negative changes in CBF, which may or may not reflect methodological confounds of the imaging modality used. In our study using PC-MRI, one individual (5% of participants) exhibited a decrease in CBF in session 1, five individuals (24%) exhibited a decrease in session 2, and two individuals (10%) exhibited a decrease in session 3, highlighted in white in Figure 2. One participant, Subject 7, showed a decrease in CBF in response to hypoxia in all three sessions, while all other participants with instances of negative hypoxic CBF changes only showed decreases in one or two sessions (Supplemental Table 1). Furthermore, many participants exhibited minimal CBF responses to hypoxia, with nineteen instances of nCBF changes of less than 2 mL/100g/min (the median absolute change between back-to-back PC acquisitions in steady state, shown in Supplemental Figure 2). Only two instances of nCBF changes of less than 2 mL/100g/min were recorded in hypercapnia, both in Subject 7 (who also showed a consistent paradoxical response to hypoxia, described above).

### Hypoxic CVR and Concurrent Changes in P_ET_CO_2_

While resting levels of P_ET_CO_2_ were targeted during the hypoxic experimental phase, P_ET_CO_2_ was on average 0.4±0.9 mmHg lower during hypoxia than during baseline. During data processing, it was noted that these unintentional concurrent changes in P_ET_CO_2_, while minor, were moderately but significantly correlated (r=0.4, p<0.01) with hypoxic CBF changes (Fig. 4). Based on this relationship, we implemented a novel correction strategy to account for such changes in P_ET_CO_2_. The subject- and session-specific hypercapnic CVR measurement was used to estimate the CBF change attributable to the unintentional change in P_ET_CO_2_ during hypoxia, and hypoxic CVR was re-calculated with this estimated CBF contribution removed.

**Figure 4.**
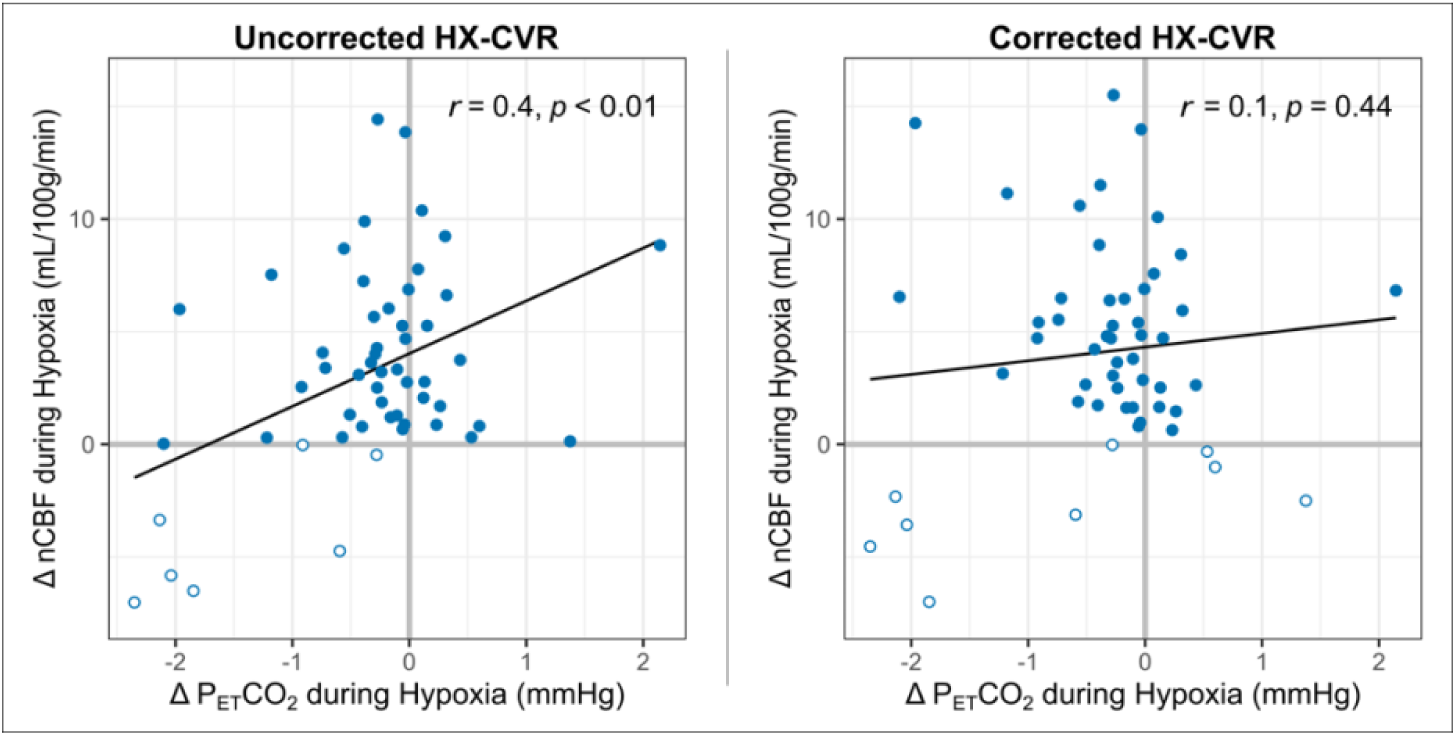
Relationship between Changes in P_ET_CO_2_ and nCBF during Hypoxia. The correlations between nCBF changes in response to hypoxia and concurrent changes in P_ET_CO_2_ are shown before (left) and after (right) correction for such P_ET_CO_2_ changes. Pearson’s correlation coefficients and p-values are included. Decreases in blood flow are indicated with open circles; all available data are shown.

P_ET_CO_2_ correction slightly increased the average nCBF change during hypoxia to 4±5 mL/100g/min or 6±7%, and average hypoxic CVR increased to 0.12±0.16 %/-mmHg P_ET_O_2_ and 0.82±1.04 %/-%SpO_2_. The correlation between concurrent P_ET_CO_2_ changes and hypoxic CBF changes became weaker and non-significant (r=0.1, p=0.44), as shown in Figure 4. Removing potential outliers did not change this interpretation (Supplemental Figure 4). With correction for P_ET_CO_2_ changes during hypoxia, three further instances of CBF decreases were identified in session 1; only one case of a negative CBF change became positive with this correction (Supplemental Figure 5). Inter-subject SDs for P_ET_O_2_ and SpO_2_ hypoxic CVR changed to 0.07%/-mmHg P_ET_O_2_ and 0.40%/-%SpO_2_, respectively, while inter-session SDs increased to 0.15%/-mmHg P_ET_O_2_ and 1.02%/- %SpO_2_, respectively, indicating decreased inter-subject but increased inter-session variability. As such, ICCs ultimately *decreased* to 0.37 and 0.31 for P_ET_O_2_ and SpO_2_ hypoxic CVR, respectively, as further instances of negative CVR were revealed. The impact of this correction on reproducibility is shown graphically in Supplemental Figure 6, and further details on the implementation of the correction method are included in the Supplemental Material.

### Hypoxic CVR and Hypercapnic CVR

Overall, hypercapnic CVR and hypoxic CVR demonstrated a positive but weak relationship, shown in Figure 5. Hypercapnic CVR and hypoxic P_ET_O_2_ CVR demonstrate a coefficient of determination (R^2^) of 0.32, while hypercapnic CVR and hypoxic SpO_2_ CVR demonstrate an R^2^ of 0.42. P_ET_CO_2_ correction of hypoxic CVR values increased this relationship slightly, with coefficients of determination increasing to 0.44 and 0.47 for P_ET_O_2_ and SpO_2_ respectively. The positive relationships between hypercapnic and hypoxic CVR metrics indicate that differences in overall vascular reactivity contribute to the inter-subject variance observed as expected, but substantial variance in hypoxic CVR (>50%) cannot be explained by differences in overall reactivity.

**Figure 5.**
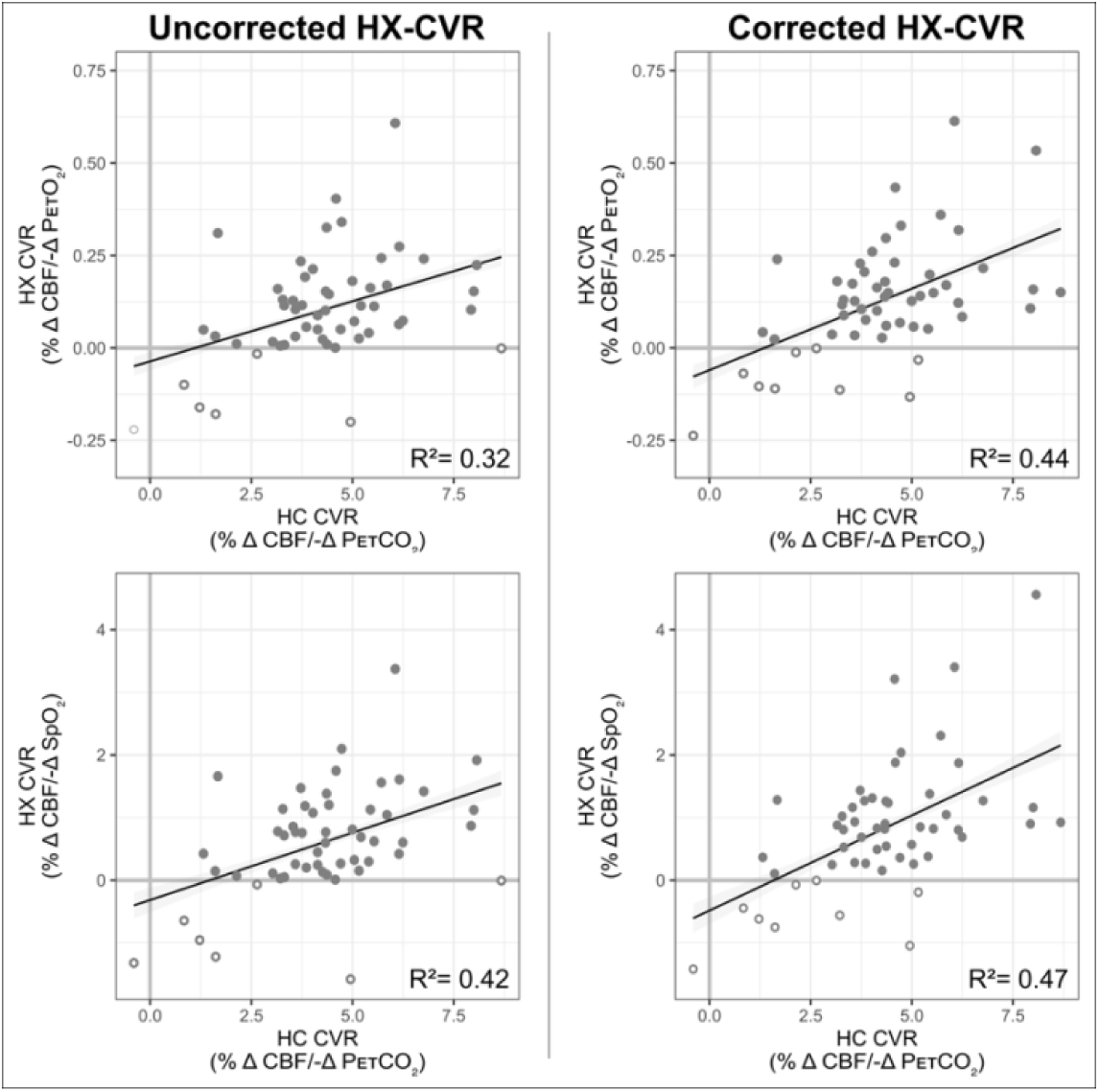
Relationship between Hypoxic and Hypercapnic CVR. The relationship between hypercapnic and uncorrected (left) and corrected (right) hypoxic CVR is shown, with hypoxic CVR relative to P_ET_O_2_ on top and hypoxic CVR relative to SpO_2_ on the bottom. Coefficients of determination, calculated from linear mixed-effects models, are included; the relationship between hypercapnic and hypoxic CVR was significant (p less than a threshold 0.05/4 = 0.0125) for all four models. Negative CVR values (both hypoxic and hypercapnic) are indicated with open circles; data from all subjects and sessions where hypoxic and hypercapnic CVR were available are included.

## Discussion

This study quantified the CBF response to moderate hypoxia and mild hypercapnia using PC-MRI and assessed CVR to each stimulus. CVR was measured in three sessions, separated by approximately three weeks and four months. While previous studies have investigated the blood flow response to hypoxia^14,21,25–28,33^, this is the first study to our knowledge to do so with PC-MRI and to assess variability in this metric. In particular, we aimed to investigate whether previously-observed inter-subject variability in hypoxic blood flow responses stems from methodological confounds, day-to-day changes, or characteristics inherent to certain individuals.

CBF is known to increase in response to hypercapnia and to mild-to-moderate hypoxia to maintain sufficient supply of nutrients and oxygen to cerebral tissue^33^. Overall, we observed blood flow responses to both stimuli in agreement with this understanding, demonstrating a small increase during hypoxia and a more substantial increase during hypercapnia. CBF increased 5±7% during hypoxia and 22±11% during hypercapnia, corresponding to CVR metrics of 0.10±0.14%/-mmHg P_ET_O_2_ or 0.59±0.88%/-%SpO_2_ for hypoxia and 4.33±1.83%/mmHg P_ET_CO_2_ for hypercapnia. Measured hypercapnic CVR in this study is consistent with previous literature, which reports blood flow increases of 3-6% per mmHg P_ET_CO_2_ when measured with ASL or PC-MRI^10^. However, previous studies have generally reported higher CVR during hypoxia than we observe in this study: using ASL to measure perfusion changes in cerebral grey matter, Harris et al. observed an increase of 11.7 mL/100g/min, corresponding to hypoxic CVRs of approximately 0.21%/- mmHg P_ET_O_2_ and 1.02%/-%SpO_2_, while Nöth et al. saw approximately 0.11%/-mmHg P_ET_O_2_ and 0.70%/-%SpO_2_^21,28^. Measuring hypoxic CVR with H_2_^15^O positron emission tomography, Binks et al. found an average increase of 6mL/100g/min, corresponding to CVRs of approximately 0.31%/-mmHg P_ET_O_2_ and 0.89%/-%SpO_2_^14^. The greater magnitude of previously reported hypoxic CVR values, particularly with respect to P_ET_O_2_, is potentially explained by their use of more severe hypoxic stimuli than the present study. Blood flow changes during hypoxia have a curvilinear relationship with P_ET_O_2_ changes, and milder stimuli can produce smaller hypoxic CVR values^33^. Furthermore, gray matter, as assessed by Harris et al. and Nöth et al., exhibits greater CVR than global metrics, as measured in the present study^1^. Further work could better characterize the dynamics of the nonlinear hypoxic whole-brain blood flow response by stepping through graded hypoxic stimuli^11,34^ (as done extensively with hypercapnic stimuli^26,35–38)^ and by employing both whole-brain and regionally-sensitive imaging metrics.

### CVR Reliability

While no previous studies have assessed the reliability of hypoxic CVR, hypercapnic CVR has been determined to be a fairly stable metric of vascular health with good reproducibility^39,40^. CVR reflects the ability of the vasculature to dilate under stress, a characteristic that is not expected to change significantly over relatively short periods except in cases of pathology or in response to therapy^41^. In healthy individuals, hypercapnic CVR has been found to typically have ICCs of more than 0.7 between sessions and inter-session coefficients of variation (CV) of 16-26%^36,39,40,42–44^. Our study produced an overall ICC of 0.67, with an inter-session SD of 1.37%/mmHg (CV=34%). Importantly, many factors can affect the reproducibility of CVR metrics, including time of day, caffeine intake, time between scans, and imaging modality^44^. We minimized these effects by scanning at the approximate same time of day and asking participants to refrain from caffeine consumption. However, our reliability is likely decreased by the longer period between scans relative to most literature (3-5 months compared with less than one week^44^) and by the imaging modality, as blood flow responses have lower reproducibility than BOLD responses^36^.

We predicted that hypoxic CVR would show similar reliability to hypercapnic CVR. However, we found that hypoxic CVR relative to P_ET_O_2_ or SpO_2_ changes had higher variability relative to the magnitude of expected changes compared to hypercapnic CVR and overall ICCs of 0.42 and 0.56 relative to P_ET_O_2_ and SpO_2_ changes, respectively, indicating poor-to-moderate reliability^45^. The magnitude of the CBF response varied greatly day-to-day, and while the directionality of most individuals’ responses was consistent between sessions, including one participant with a consistent decrease in CBF during hypoxia, several subjects exhibited negative hypoxic CVR in only one or two sessions.

### Paradoxical CBF Responses to Hypoxia

Although blood flow is generally acknowledged to increase during hypoxia, in line with our overall average observations in this study, several previous studies of hypoxic CVR have identified unexpected variability between individuals in the cerebrovascular response to moderate hypoxic hypoxia. Nöth et al. found that 30% of participants had a *decrease* in global CBF, while Binks et al. reported that one of five subjects had no significant change in CBF^14,28^. The rate of minimal or paradoxical responses was similar to previously published literature, with 1-5 participants, or 5-24% of subjects, showing a decrease in CBF in response to hypoxia during a given scan session. Furthermore, 3-9 participants, or 16-43% of subjects, did not show a meaningful change in CBF in response to hypoxia during a given scan session. The source of this variability is unclear. Previous work has relied on assumptions of unchanged labeling efficiency and T1 of arterial blood, both of which may change with hypoxia and confound CVR estimations^28^. However, using a method unbiased by these assumptions, we still observe paradoxical responses, indicating that these responses do not stem exclusively from measurement biases.

Paradoxical CBF responses to hypoxia could be caused by overall differences in the ability of the vasculature to react or by a truly atypical cerebrovascular response to hypoxia. In the lungs, arteries constrict in response to decreased oxygen concentrations in the alveoli^46^; similar vasoconstrictive mechanisms may be involved in regions of the brain in hypoxia, leading to competing vasoconstrictive and vasodilatory effects producing the variability seen in the global CBF response^47^. Lawley et al. verified that observed reductions in regional blood flow during hypoxia was caused by vasoconstriction, rather than an inability for the vasculature to dilate, as the regions in question displayed the typical vasodilatory response to a subsequent hypercapnic challenge^48^.

Prior studies have also not investigated the reliability of hypoxic CVR measures, leaving it ambiguous whether these minimal and negative responses are characteristic of certain individuals. If this response is a reliable trait of an individual, it could have substantial implications regarding the vulnerability of their cortical tissue to damage caused by moderate hypoxia. Of note, we observed one participant (S7, Supplemental Table 1) who demonstrated negative blood flow responses to hypoxia in all three sessions. Further work with larger sample sizes will be needed to predict which individuals are likely to show a paradoxical CBF response to hypoxia.

### Sources of CVR Variability

Experimental methodology and general data quality likely still contribute to the observed variability in our PC-MRI measurements of CBF during different inhaled gas challenges. While not directly assessed in this study, motion artifacts could have impacted data quality and therefore flow quantification, particularly due to changes in ventilation induced by the respiratory stimuli. The requirement for a gas delivery face mask necessitated the use of a 32-channel coil as opposed to the 64-channel coil preferred for neck imaging, potentially resulting in a lower signal-to-noise ratio and thereby increasing flow measure variability. Subtle inconsistencies in vessel segmentation and slice placement across sessions, although minimized, may have further introduced variability in flow calculations. The implemented PC acquisition employed eight cardiac bins, resulting in a temporal resolution that may have been insufficient to fully capture the dynamics of pulsatile flow. Additionally, the fixed order of respiratory challenges may have contributed to the enhanced variability observed in hypoxic CVR compared to hypercapnic CVR. Although prioritizing hypoxic CVR ensured its collection even if participants became uncomfortable, acquiring these measurements before the hypercapnic phase may have resulted in greater physiological variability as participants were still acclimating to the MR and gas challenge environment.

Variability in the measured CBF response to hypoxia could also result if CBF was not given enough time to stabilize under the stimulus. The blood flow response to hypoxia in gray matter is delayed by more than three minutes, with an additional 4.5 minutes required to reach steady state^21^; we therefore allowed approximately 7.5 minutes before collecting steady-state CBF measures under hypoxia. We verified that blood flow had stabilized following the transition to hypoxia, as demonstrated by the differences in CBF measured in back-to-back repetitions of PC-MRI, indicating that instability of blood flow is not a dominant driver of the observed variability and select paradoxical responses.

Changes in P_ET_CO_2_ levels substantially impact CBF, and concomitant changes in P_ET_CO_2_ during hypoxia administration have been identified as a confound in isolating the hypoxic blood flow response and suggested as a potential source of variability^14,28^. As arterial carbon dioxide and oxygen concentrations are closely linked through respiration, changes in oxygen levels cause unintentional modulations of carbon dioxide concentrations, and hypoxia typically results in hypocapnia due to increased ventilation unless P_ET_CO_2_ is externally controlled. This hypocapnia can result in a competing vasoconstrictive effect, potentially obscuring the true hypoxic blood flow response. We used a computer-controlled gas blending system to clamp P_ET_CO_2_ at baseline levels. However, a slight decrease in P_ET_CO_2_ during hypoxia was still observed, and individual changes in P_ET_CO_2_ during hypoxia were found to be correlated with hypoxic nCBF changes (Figure 4). We therefore corrected CBF measurements during hypoxia for these effects, estimating the impact of the change in P_ET_CO_2_ from subject- and session-specific hypercapnic CVR measures.

This correction removed the significant impact of confounding P_ET_CO_2_ changes on measures of hypoxic CVR (Fig. 4); however, it revealed further instances of decreased blood flow in response to hypoxia, and ICCs were lower following correction. This may mean that our correction approach makes us more sensitive to true physiologic variability in hypoxic CVR across sessions, although we cannot conclude this using our current study data. The reductions in reliability suggest, however, that the observed inter-session variability in hypoxic CBF responses likely does not stem from concurrent P_ET_CO_2_ changes.

Variability in the hypoxic CBF response may instead result from variability in the degree of hypoxia achieved. Targets for P_ET_O_2_ were selected for each subject in a practice session outside the scanner and were chosen to maintain participants at a SpO_2_ of 85%. However, many participants demonstrated smaller changes in oxygenation during MRI sessions despite identical P_ET_O_2_ targets and substantial variability between scan sessions (shown in Supplemental Figure 7), and the average SpO_2_ during hypoxia was only 90%. While previous literature indicates that there exists a linear relationship between arterial deoxygenation and CBF changes in hypoxia^33^, some studies have suggested that the arterial oxygenation threshold for a blood flow response to hypoxia is 90%^49^. Given this, we would expect participants who did not achieve SpO_2_ values below 90% to also not show meaningful increases in CBF (<2mL/100g/min) during hypoxia, and participants who achieve SpO_2_ values below this 90% threshold to demonstrate substantial CBF increases. This scenario is indicated by shading in Figure 6. Contrary to these literature predictions, many subjects in the present study demonstrated an increase in CBF at a SpO_2_ of greater than 90% or insubstantial changes in CBF with SpO_2_ levels well below this threshold, and still others displaying *decreased* CBF despite dramatic decreases in SpO_2_. While it is likely that we were unable to reach individual thresholds for vasoactive responses in some subjects, the lack of a strong relationship between CBF changes in hypoxia and SpO_2_ decreases (r=0.21 uncorrected, r=0.11 corrected for concurrent P_ET_CO_2_ changes) suggests that the paradoxical or minimal responses we observed are not all attributable to insufficient decreases in oxygenation and may instead be due to other sources. Still, a greater degree of hypoxia may decrease inter- and intra-subject variability in hypoxic blood flow responses, and future studies should investigate whether a greater degree of hypoxia improves the reliability of hypoxic CVR measures.

**Figure 6.**
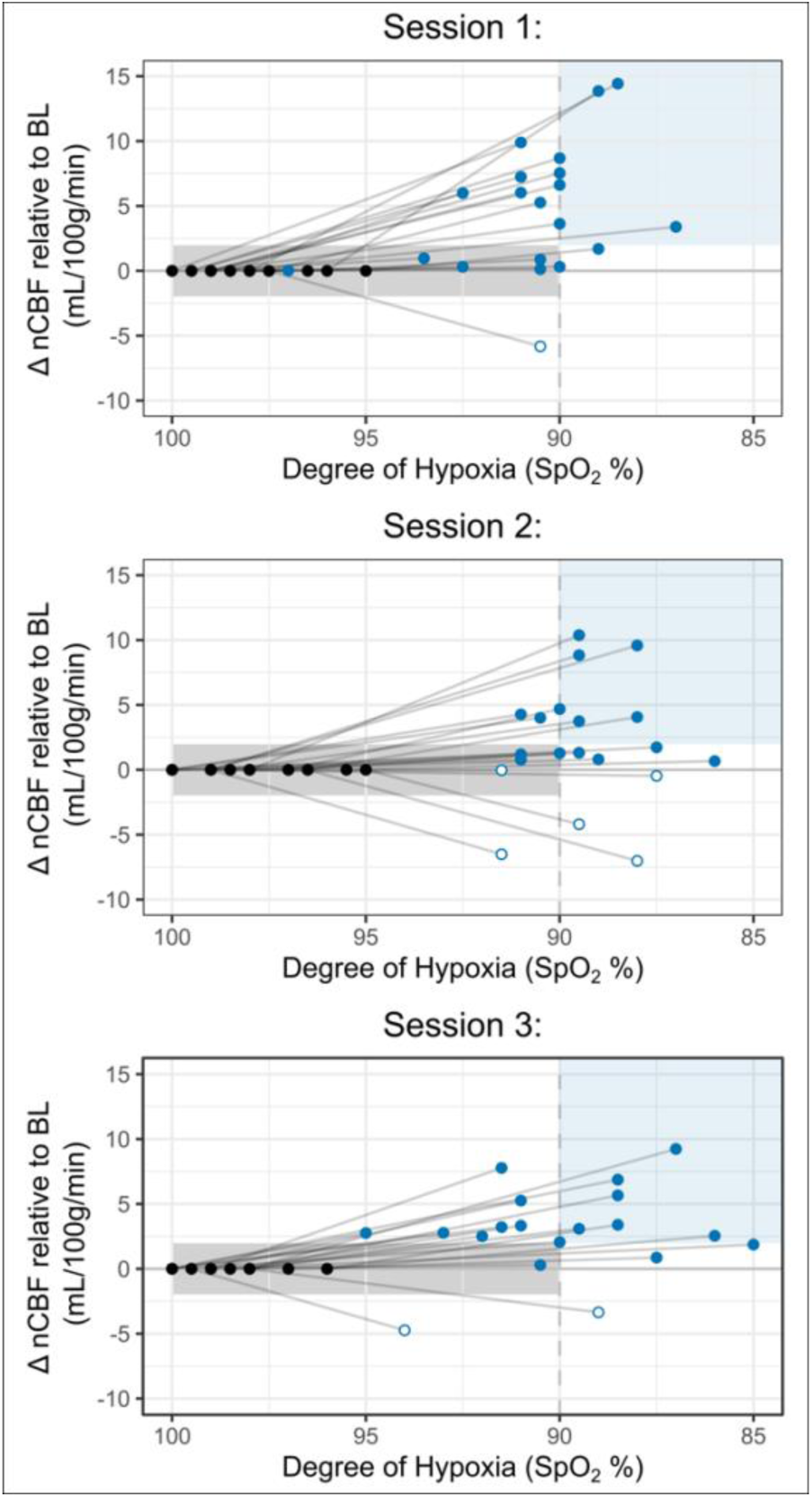
Comparisons of nCBF changes with SpO_2_ Decreases during Hypoxia. Changes in nCBF relative to baseline are shown during hypoxia (blue) and baseline (black) relative to the degree of hypoxia, measured by SpO_2_. Approximate ranges for expected changes in nCBF relative to SpO_2_, based on previous literature^49^, are shaded in light grey (greater than 90% SpO_2_) and light blue (less than 90% SpO_2_). Open circles indicate negative nCBF changes; all available data are included.

### Overall Vascular Reactivity and Distinct Effects of Hypoxia and Hypercapnia

We compared hypercapnic and hypoxic measures of CVR to assess whether observed variability may be attributable to differences in overall cerebrovascular responsiveness. Across all sessions, less than half of variance is shared between hypercapnic and hypoxic reactivity, suggesting that additional sources of variability are present in the vascular response to hypoxia that are not present in the response to hypercapnia. The observed variance in hypoxic cerebral blood flow responses warrants further investigation, as hypoxic CVR may provide distinct information about vascular health not revealed by hypercapnic CVR.

In conclusion, we investigated hypoxic CVR alongside hypercapnic CVR using PC-MRI and mild-to-moderate stimuli in three sessions. We observed great variability in hypoxia responses between subjects, including paradoxical decreases in CBF in some individuals, and hypoxic CVR metrics showed poor-to-moderate reliability in comparison with hypercapnic CVR metrics, which demonstrated moderate reliability^45^. Methodological confounds did not account for the high variability and paradoxical responses observed; PC-MRI is free of the assumptions that may bias CVR measures made with BOLD or ASL MRI, and neither concurrent changes in P_ET_CO_2_ nor variability in deoxygenation achieved explained the noted variability in hypoxic CVR. While hypoxic and hypercapnic CVR demonstrated a positive relationship, less than half of their variance was shared, indicating that hypoxic CVR is subject to other sources of variability. We show that while paradoxical instances of hypoxic CVR likely reflect true inter-subject variations in the cerebrovascular response, further investigation of hypoxic CVR is warranted to understand the sources of this variability and its implications for cerebrovascular health.

## Acknowledgements

This work was supported by the Center for Translational Imaging at Northwestern University. The authors would like to thank Andrew Vigotsky for his statistical guidance and Rachael Young for her assistance in data collection.

## Author Contribution Statement

**Hannah R Johnson**: Investigation, Formal analysis, Project administration, Visualization, Writing – original draft, Writing – review & editing. **Max C Wang**: Investigation, Writing – review & editing. **Rachael C Stickland**: Conceptualization, Methodology, Investigation, Writing – review & editing. **Yufen Chen**: Conceptualization, Methodology, Writing – review & editing, Funding Acquisition. **Todd Parrish**: Conceptualization, Methodology, Writing – review & editing, Funding Acquisition. **Farzaneh A Sorond**: Conceptualization, Methodology, Writing – review & editing, Funding Acquisition. **Molly G Bright**: Conceptualization, Methodology, Project Administration, Writing – review & editing, Supervision, Funding Acquisition.

## Supplementary Material

Supplemental material for this article is available online.

