## Supplemental material for "Variable Cerebral Blood Flow Responsiveness to Acute Hypoxic Hypoxia"

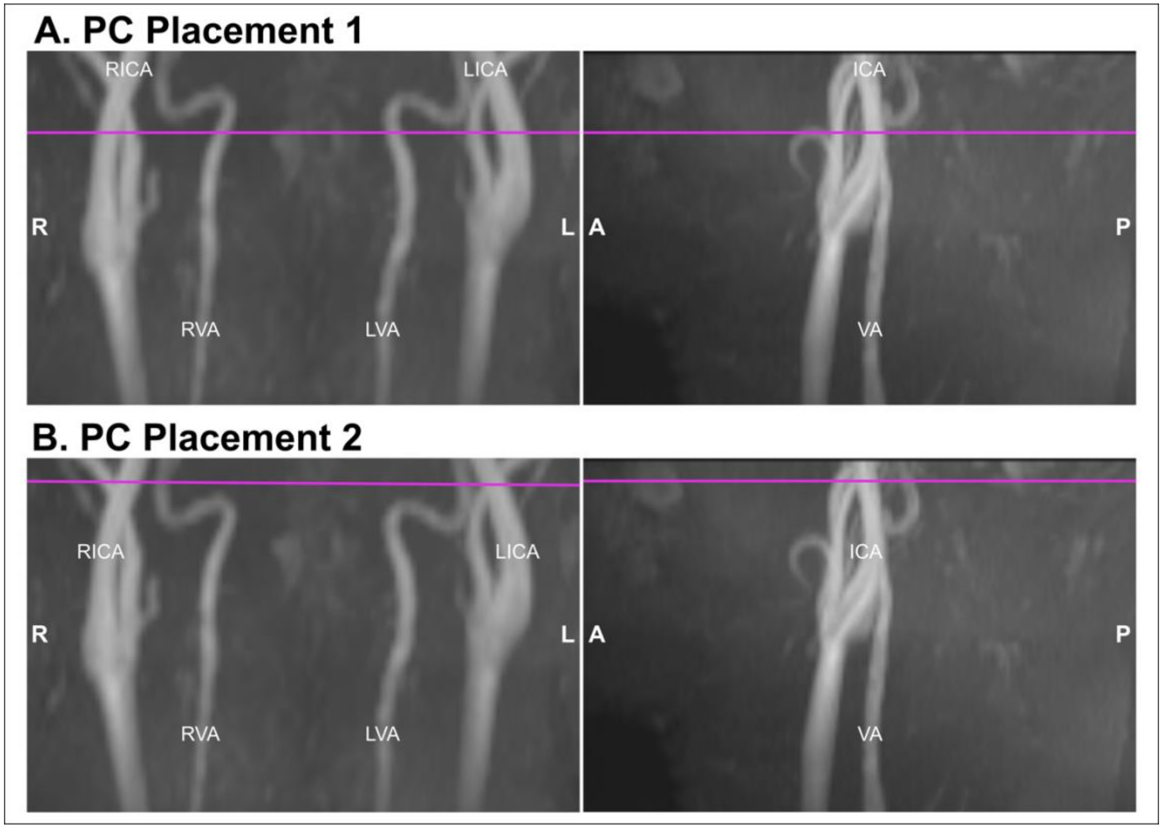


***Supplemental Figure 1. Example Slice Placements.*** *Placement of PC imaging slices for an example participant are shown in purple, overlaid on a 3D time-of-flight angiogram. Placement 1 (top) prioritized orthogonality with the internal carotid arteries and was collected twice during steady-state in each respiratory phase. Placement 2 (bottom) prioritized orthogonality with any vessel(s) that was not considered orthogonal in Placement 1, particularly vertebral arteries, and was collected once in each respiratory phase. In this example participant, both vertebral arteries are not optimally orthogonal in Placement 1, and are therefore better captured in Placement 2. Right (R) and left (L) internal carotid arteries (ICA) and vertebral arteries (VA) are labeled.x*

| | Hypoxia | | | | | | $P_{ET}CO_2$-Corrected Hypoxia | | | | | | Hypercapnia | | |
|---|---|---|---|---|---|---|---|---|---|---|---|---|---|---|---|
| | S1 | | S2 | | S3 | | S1 | | S2 | | S3 | | S1 | S2 | S3 |
| Subject | $P_{ET}O_2$ CVR | $SpO_2$ CVR | $P_{ET}O_2$ CVR | $SpO_2$ CVR | $P_{ET}O_2$ CVR | $SpO_2$ CVR | $P_{ET}O_2$ CVR | $SpO_2$ CVR | $P_{ET}O_2$ CVR | $SpO_2$ CVR | $P_{ET}O_2$ CVR | $SpO_2$ CVR | $P_{ET}CO_2$ CVR | $P_{ET}CO_2$ CVR | $P_{ET}CO_2$ CVR |
| 1 | 0.05 | -- | 0.06 | -- | 0.17 | 1.04 | -0.04 | -- | -0.02 | -- | 0.17 | 1.05 | 4.57 | 5.52 | 5.85 |
| 2*† | 0.40 | 1.75 | -0.16 | -0.96 | 0.07 | 0.32 | 0.43 | 1.88 | -0.10 | -0.62 | 0.06 | 0.26 | 4.59 | 1.22 | 5.04 |
| 3*† | 0.61 | 3.37 | 0.13 | 0.86 | 0.09 | 0.45 | 0.61 | 3.40 | 0.17 | 1.16 | 0.16 | 0.83 | 6.05 | 3.53 | 4.14 |
| 4 | 0.04 | 0.27 | 0.00 | -0.01 | 0.06 | 0.20 | -- | -- | 0.15 | 0.92 | 0.08 | 0.27 | -- | 8.68 | 3.86 |
| 5*† | 0.00 | 0.01 | -0.02 | -0.07 | 0.33 | 1.38 | 0.23 | 3.21 | 0.00 | 0.00 | 0.30 | 1.26 | 4.57 | 2.64 | 4.36 |
| 6*† | 0.27 | 1.61 | 0.15 | 0.77 | 0.11 | 0.71 | 0.32 | 1.87 | 0.18 | 0.90 | 0.13 | 0.81 | 6.16 | 4.34 | 3.31 |
| 7*† | -0.18 | -1.22 | -0.22 | -1.32 | -0.10 | -0.64 | -0.11 | -0.75 | -0.24 | -1.42 | -0.07 | -0.45 | 1.62 | -0.40 | 0.84 |
| 8*† | 0.19 | 1.19 | 0.34 | 2.10 | 0.23 | 1.47 | 0.21 | 1.27 | 0.33 | 2.04 | 0.23 | 1.44 | 3.83 | 4.73 | 3.72 |
| 9 | 0.21 | 1.08 | 0.04 | 0.18 | -- | -- | 0.26 | 1.31 | -- | -- | -- | -- | 4.02 | -- | -- |
| 10 | 0.05 | 0.42 | -0.19 | -1.30 | -0.20 | -1.58 | 0.04 | 0.37 | -- | -- | -0.13 | -1.05 | 1.32 | -- | 4.95 |
| 11† | 0.03 | 0.26 | 0.15 | 1.12 | 0.12 | 0.75 | 0.03 | 0.28 | 0.16 | 1.16 | -- | -- | 3.59 | 8.00 | -- |
| 12*† | 0.13 | 1.14 | 0.31 | 1.66 | 0.03 | 0.15 | 0.12 | 1.02 | 0.24 | 1.28 | 0.02 | 0.11 | 3.28 | 1.67 | 1.61 |
| 13*† | 0.16 | 1.13 | 0.11 | 0.69 | 0.07 | 0.60 | 0.20 | 1.38 | 0.14 | 0.85 | 0.08 | 0.69 | 5.44 | 5.21 | 6.24 |
| 14*† | 0.22 | 1.92 | 0.03 | 0.15 | 0.10 | 0.60 | 0.53 | 4.56 | -0.03 | -0.19 | 0.14 | 0.82 | 8.07 | 5.16 | 4.33 |
| 15*† | 0.01 | 0.09 | 0.05 | 0.24 | 0.10 | 0.87 | 0.06 | 0.54 | 0.10 | 0.49 | 0.11 | 0.90 | 4.37 | 4.14 | 7.93 |
| 16† | 0.24 | 1.42 | 0.05 | 0.27 | -- | -- | 0.22 | 1.27 | 0.07 | 0.36 | -- | -- | 6.76 | 4.71 | -- |
| 17 | 0.24 | 1.56 | 0.21 | 1.42 | 0.15 | 1.20 | 0.36 | 2.31 | -- | -- | 0.15 | 1.23 | 5.71 | -- | 4.42 |
| 18*† | 0.01 | 0.07 | 0.02 | 0.13 | 0.12 | 0.76 | -0.01 | -0.07 | 0.03 | 0.16 | 0.11 | 0.69 | 2.14 | 4.26 | 3.76 |
| 19*† | 0.01 | 0.03 | 0.18 | 0.81 | 0.10 | 0.77 | -0.11 | -0.56 | 0.13 | 0.57 | 0.13 | 0.93 | 3.22 | 5.00 | 3.59 |
| 20*† | 0.06 | 0.42 | 0.04 | 0.30 | 0.01 | 0.05 | 0.12 | 0.80 | 0.05 | 0.38 | 0.09 | 0.52 | 6.15 | 5.40 | 3.32 |
| 21*† | 0.11 | 0.62 | 0.02 | 0.11 | 0.16 | 0.78 | 0.15 | 0.82 | 0.04 | 0.25 | 0.18 | 0.88 | 5.53 | 3.03 | 3.15 |

***Supplemental Table 1. Individual Subject CVR Values and Inclusions.*** *Cerebrovascular reactivity values for each subject and session (S1, S2, S3) are shown for hypoxia before and after corrections for unintentional changes in end-tidal carbon dioxide levels ($P_{ET}CO_2$) relative to end-tidal oxygen ($P_{ET}O_2$) (%ΔCBF/-Δ $P_{ET}O_2$) and oxygen saturation ($SpO_2$) (%ΔCBF/-Δ $SpO_2$) and for hypercapnia (%ΔCBF/-Δ $P_{ET}CO_2$). Subjects included in calculations of repeatability metrics, Group A, are indicated by asterisks; subjects included in supplemental figure 4 (Group B) are indicated by daggers.*

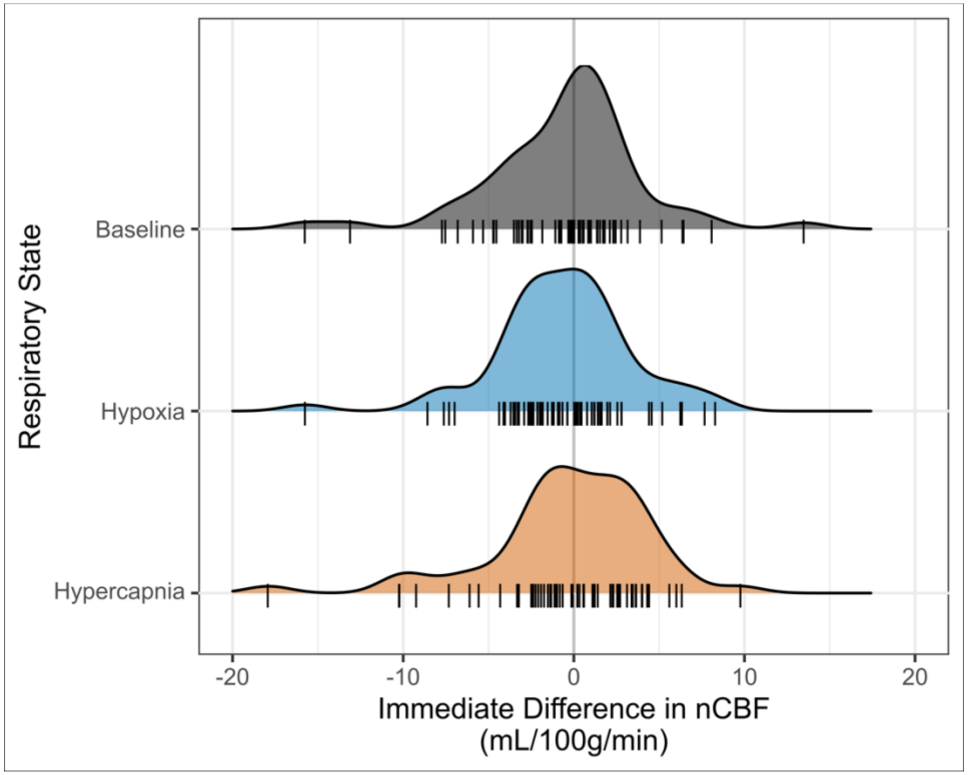


***Supplemental Figure 2. Change in nCBF Measured in Back-to-Back PC Acquisitions.*** *Immediate changes in normalized cerebral blood flow (nCBF), represented by the difference in nCBF measured in back-to-back acquisitions of phase-contrast data, are shown for each respiratory state. The baseline measurements provide an estimate of the inherent noise and error present in our data, and the insignificant changes in this metric in hypoxia (p=0.75) and hypercapnia (p=0.72) compared to baseline indicate that steady-state blood flow was reached and that no additional sources of significant noise were present during these stimuli. Across all states, the mean of the absolute value of immediate differences is 3mL/100g/min and the median is 2mL/100g/min. Rug plots, shown in black at the bottom of each respiratory state, indicate individual data points.*

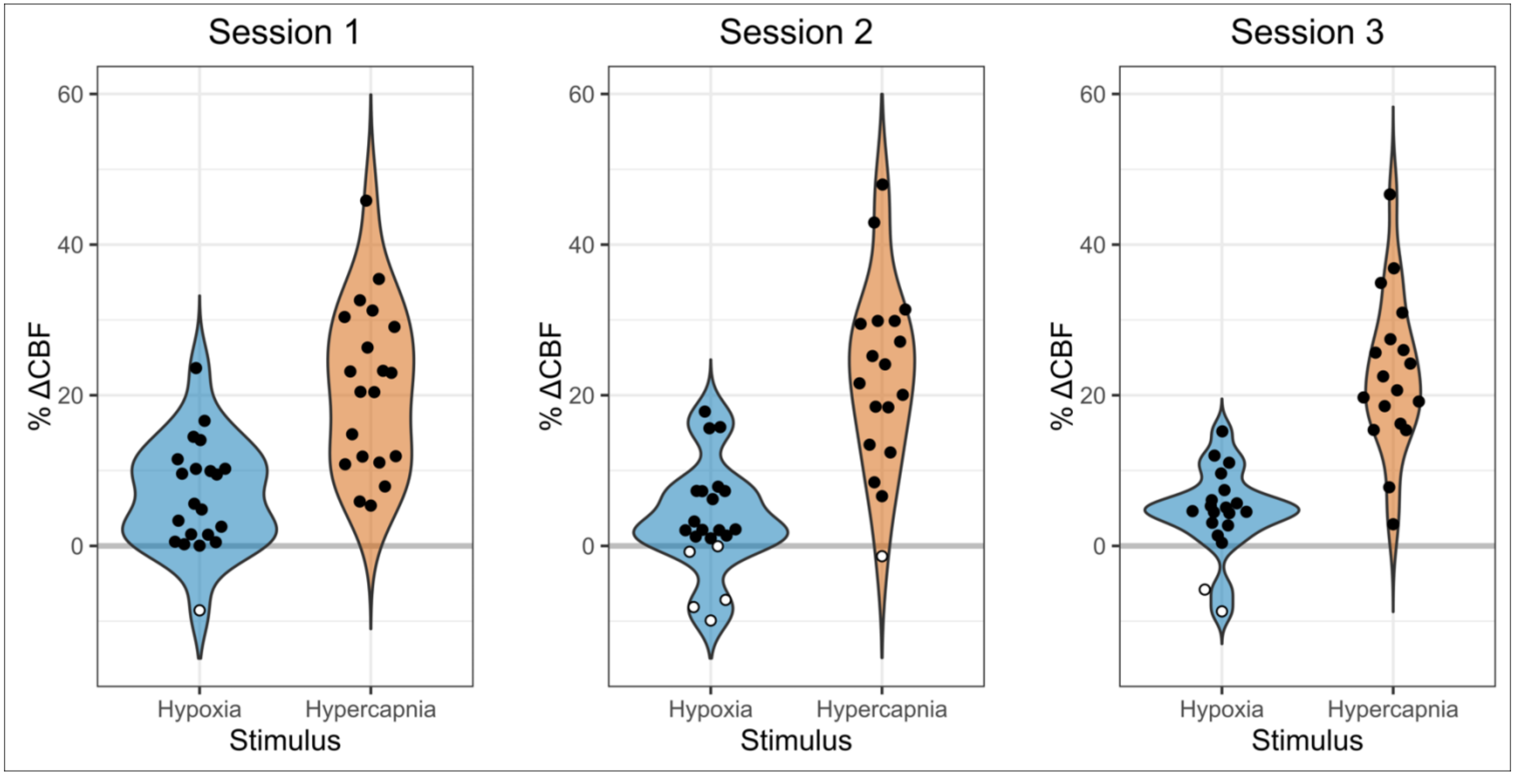


***Supplemental Figure 3. Relative Changes in Blood Flow During Hypoxia and Hypercapnia.*** *Percent change in CBF during hypoxia (blue) and hypercapnia (orange), relative to baseline, are shown for each session. Positive values, indicating an increase in CBF, are shown with black circles, while negative values, indicating a decrease in CBF, are shown with white circles. Note that sessions and gas stimuli have different numbers of subjects. Distributions vary slightly from Figure 2 due to scaling by baseline blood flow.*

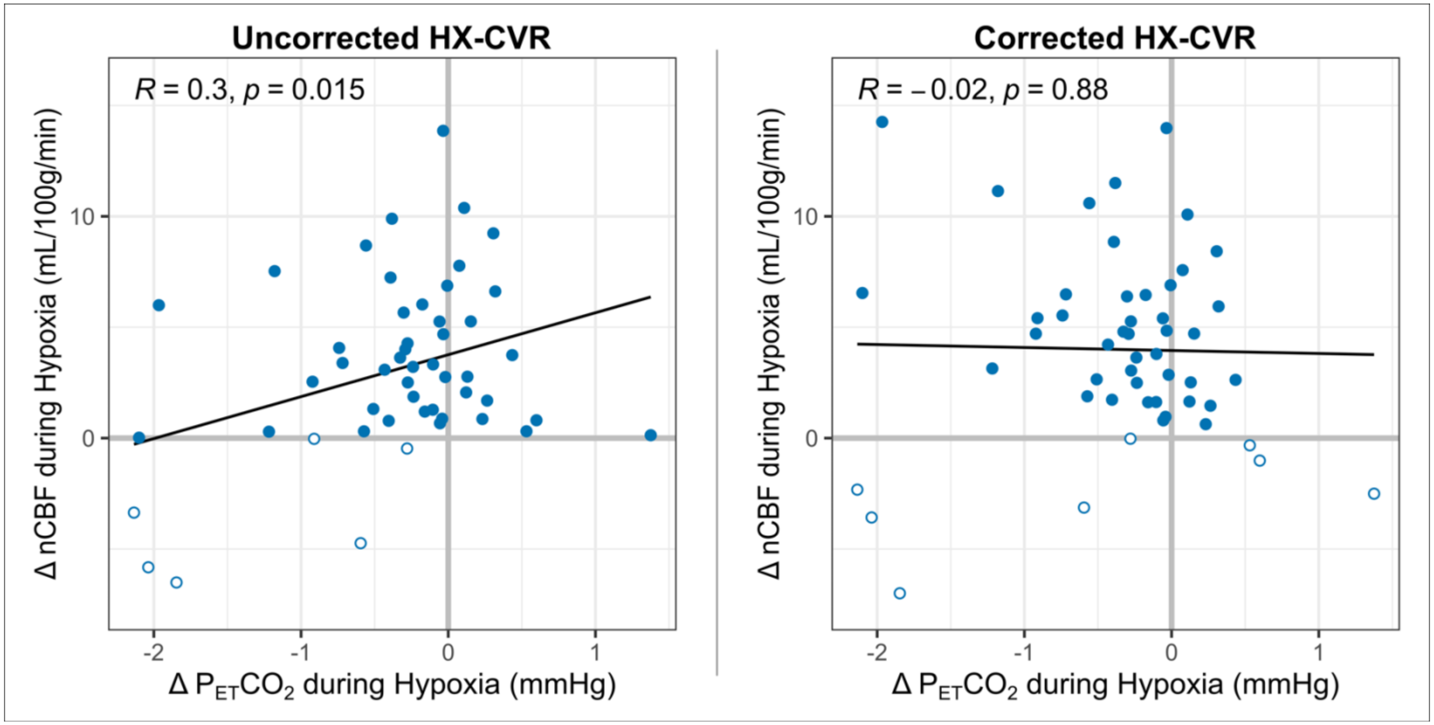


***Supplemental Figure 4. Relationship between Changes in $P_{ET}CO_2$ and nCBF during Hypoxia with Outliers Removed.*** *The correlations between nCBF changes and concurrent changes in $P_{ET}CO_2$ during hypoxia are shown before (left) and after (right) correction for such PETCO2 changes. Three outliers were identified, falling below the 1st percentile or above the 99th percentile in the distributions of $P_{ET}CO_2$ and nCBF changes: one change in PETCO2 (=-2.35mmHg) was less than the lower bound for PETCO2 changes, one change in PETCO2 (=2.14mmHg) was greater than the upper bound for PETCO2 changes, and one change in nCBF (=14.1mL/100g/min) was greater than the upper bound for nCBF changes. Pearson's correlation coefficients and p-values, calculated without outliers, are included. Decreases in blood flow are indicated with open circles; all other available data are shown.*

*Hypoxic CVR Corrections for Concurrent Changes in $P_{ET}CO_2$* For participant and session, the change in $P_{ET}CO_2$ measured during the steady-state acquisitions of CBF during hypoxia was calculated (Eq. 4). This change was multiplied by the session-specific hypercapnic CVR to obtain the correction factor (Eq. 5). This correction factor was subtracted from the uncorrected hypoxic cerebrovascular blood flow measurement to obtain a corrected measure of hypoxic CBF (Eq. 6), which was then used to calculate corrected hypoxic CVR.

$$(\Delta P_{ET}CO_2)_{HX} = (P_{ET}CO_2)_{HX} - (P_{ET}CO_2)_{BL} \quad \text{(Eq. 4)}$$

$$CF_{HX} = (\Delta P_{ET}CO_2)_{HX} * (CVR)_{HC} * (CBF)_{BL} \quad \text{(Eq. 5)}$$

$$CBF_{HX,corr} = CBF_{HX,uncorr} - CF_{HX} \quad \text{(Eq. 6)}$$

The relationships in hypoxic CVR acquired in different sessions, before and after correction for $P_{ET}CO_2$ changes, are shown in Supplemental Figure 6.

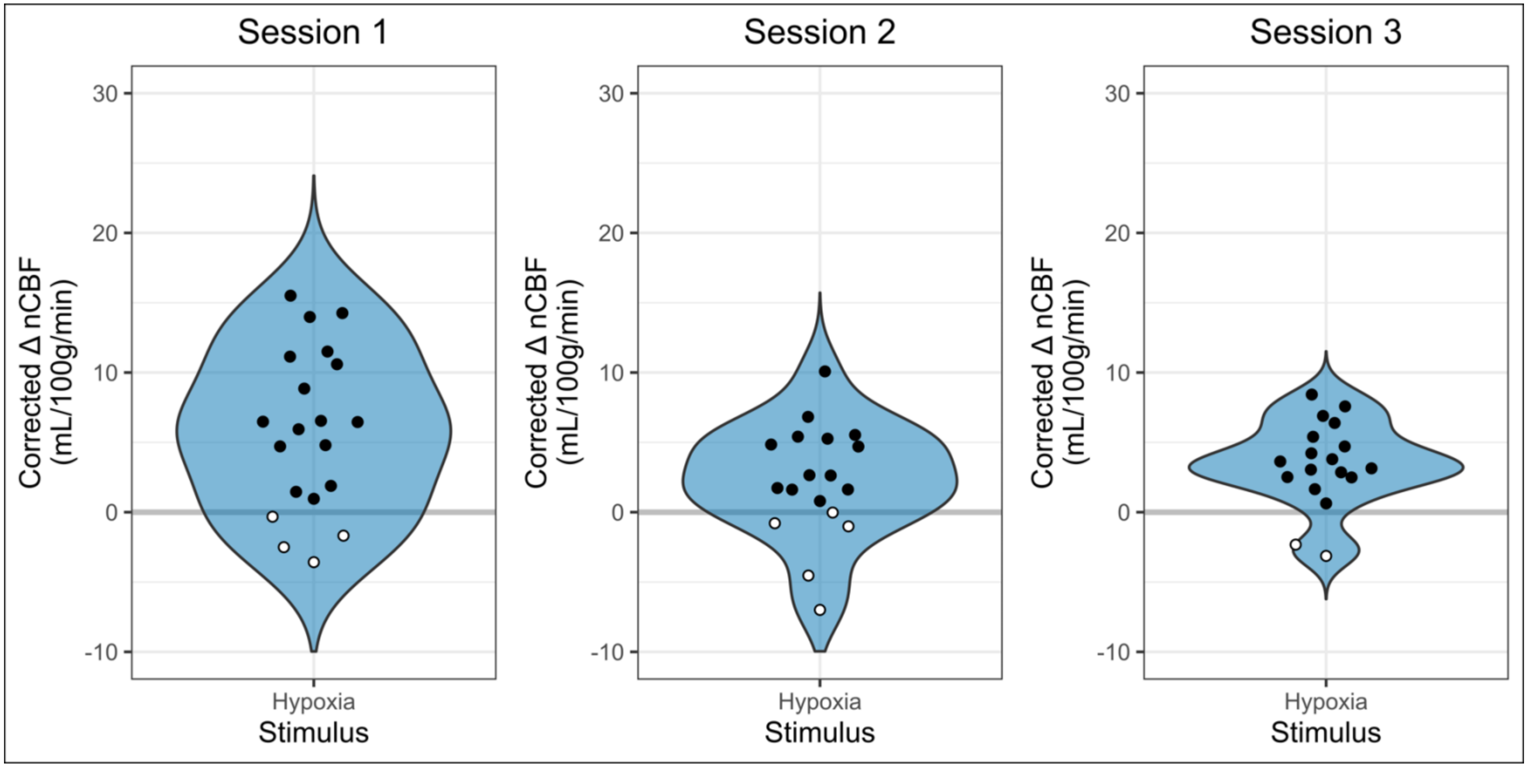


***Supplemental Figure 5. Corrected Relative Changes in CBF during Hypoxia.*** *Distributions of nCBF changes during hypoxia relative to baseline, following correction for concurrent changes in $P_{ET}CO_2$, are shown for each session (Figure 2 displays uncorrected results). Open circles indicate negative values. All subjects and sessions for which a hypercapnic CVR value was available are included.*

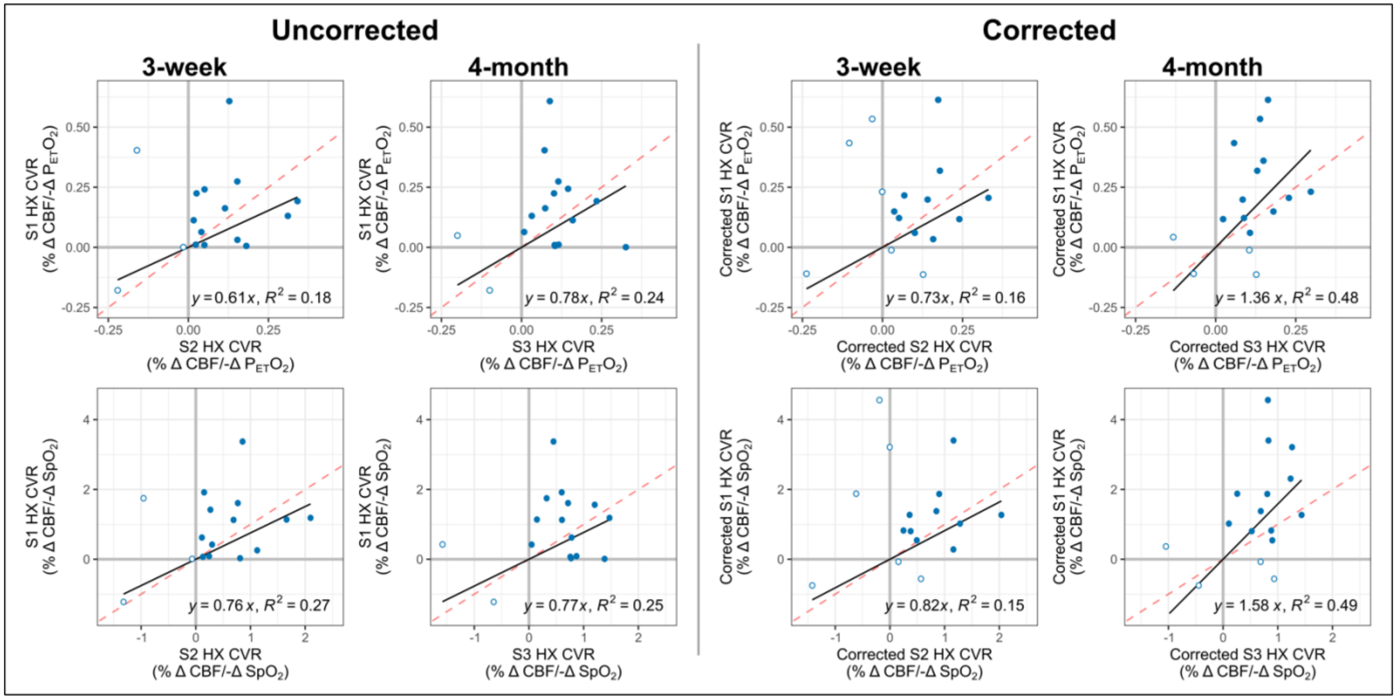


***Supplemental Figure 6. Impact of $P_{ET}CO_2$ Correction on Hypoxic CVR.*** *The correlation of hypoxic CVR metrics between scan sessions before (left) and after (right) corrections for unintentional changes in $P_{ET}CO_2$ is shown for Group B. Slopes of the line of best fit and $R^2$ values are included. The identity line, representing a slope of 1 and therefore perfect reproducibility, is plotted in red. Open circles represent negative CVR values.*

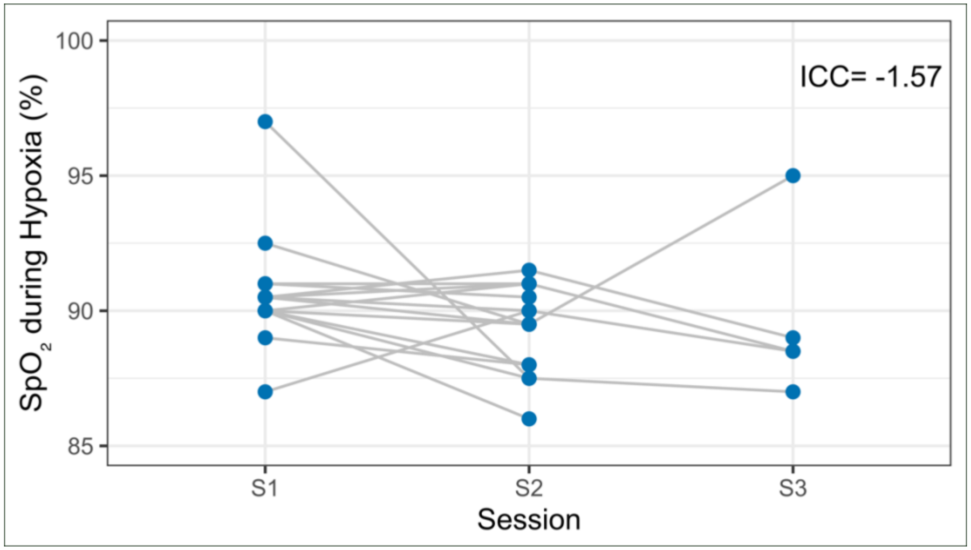


***Supplemental Figure 7. Reliability of $SpO_2$ during hypoxia.*** *The peripheral arterial oxygen saturation ($SpO_2$) value achieved during hypoxia is shown across sessions for similar achieved end-tidal oxygen ($P_{ET}O_2$) levels. Only participants and sessions for whom $P_{ET}O_2$ during hypoxia was within 1mmHg of the achieved $P_{ET}O_2$ in session 1 are included. The overall intraclass correlation coefficient (ICC) is listed for the included data; the negative value of the ICC indicates that within-subject variation exceeds between-subject variation.*
